# ASOCompass: Context- and Chemistry-Aware Activity Prediction for Transferable Antisense Oligonucleotide Screening

**DOI:** 10.64898/2026.08.03.742461

**Authors:** Shuyu Liu, Jin Zhuo, Shuchang Lei, Tianhao Wu, Jiaxuan Han, Chaoyi Wu, Yanfeng Wang, Weidi Xie

**Affiliations:** Shanghai Jiao Tong University, Shanghai, China; OneX Intelligence, Shanghai, China; University of Toronto, Toronto, ON, Canada

## Abstract

Antisense oligonucleotide (ASO) activity is jointly influenced by nucleotide sequence, chemical modification, target-RNA context, dose, delivery protocol, and cellular environment. Most existing computational screening methods model only a subset of these factors, limiting their ability to predict experimentally measured activity across heterogeneous screening conditions and previously unseen biological contexts. We introduce **ASOCompass**, a context-and chemistry-aware framework for ASO activity prediction and candidate ranking. ASOCompass integrates contextualized ASO and target-RNA sequence representations with position-specific molecular representations of chemical modifications. It further incorporates dose and delivery information together with prototype-adapted transcriptomic representations of target genes and cell lines. To encourage chemically and biophysically informative representations, the model is jointly trained on auxiliary molecular-property and sequence-derived thermodynamic prediction tasks. We evaluate ASOCompass on ASO Atlas, a large patent-derived dataset of RNase H-mediated gapmer ASOs, under held-out drug, target-gene, cell line, and joint gene–cell line settings. ASOCompass achieves an overall Spearman correlation of 0.5970, improving over the strongest ASO-specific baseline by 0.0421, and consistently performs best across all four distribution shifts. When adapted to unseen SOD1 and KLKB1 targets, ASOCompass also provides more accurate candidate ranking across different annotation budgets, reaching correlations of 0.830 and 0.696 with 1,024 target-specific labels. Additional analyses suggest that molecular-property supervision improves modification-specific ranking, while the auxiliary thermodynamic task produces representations more closely aligned with measured inhibition. These results demonstrate the potential of jointly modeling sequence, chemistry, and experimental-biological context for transferable ASO screening.

## Introduction

**Antisense oligonucleotides (ASOs)** are programmable, single-stranded nucleic acid therapeutics that modulate RNA through sequence-specific hybridization, enabling RNA degradation, splice modulation, or steric blocking (Crooke et al. 2021b; Liu et al. 2025). By targeting disease-associated RNAs, ASOs offer therapeutic opportunities for genetic disorders poorly addressed by small molecules or proteins (Crooke et al. 2021a; Tambuyzer et al. 2020). Despite their therapeutic promise, the selection of active ASO candidates remains experimentally intensive because their activity is not determined by sequence complementarity alone (Bennett 2019). As shown in Figure 1, ASO activity is jointly governed by nucleotide sequence, which determines target recognition, local hybridization, and potential off-target interactions (Hagedorn et al. 2017; Ho et al. 1998); sugar and backbone modifications, which regulate binding affinity, nuclease stability, cellular uptake, and toxicity (Shen and Corey 2018; Khvorova and Watts 2017); and experimental and biological context, including dose, delivery protocol, target abundance, and cellular state (Roberts, Langer, and Wood 2020; Juliano 2016). Their interactions can produce substantially different inhibition for the same nucleotide sequence across chemical patterns and evaluation contexts, creating a combinatorial design space difficult to explore through iterative synthesis and wet-lab screening (Linnane et al. 2019; Mor, Avkin-Nachum, and Dominissini 2025).

**Figure 1:**
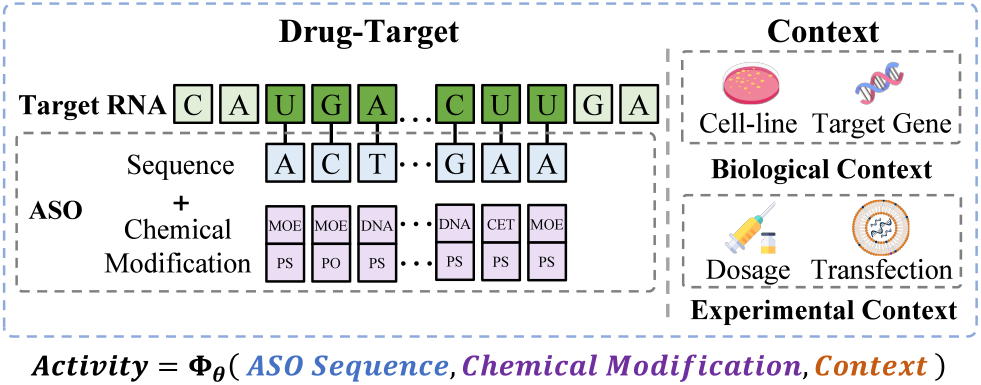
Major determinants of ASO activity. ASO activity is jointly determined by nucleotide sequence, chemical modification, and the biological and experimental contexts.

Computational ASO design has progressed from rule-based scoring of sequence composition (Krieg, Guga, and Stec 2003), target accessibility (Andersson et al. 2025; Ding, Chan, and Lawrence 2004), and hybridization energy (Lorenz et al. 2011; Mathews et al. 1999) to data-driven activity prediction (Hwang et al. 2024, 2026). Recent models have incorporated neural sequence encoders, local target-RNA contexts, chemical-modification categories, and available experimental metadata (Hill et al. 2026; Lv et al. 2026),demonstrating the value of learning directly from measured inhibition data. Nevertheless, robust prediction across new ASO designs, target genes, and cellular environments remains challenging, suggesting that current representations do not fully capture the factors governing experimentally observed activity (Bennett and Swayze 2010; Shadid, Badawi, and Abulrob 2021).

Three limitations are particularly important: (i) existing models provide only limited representations of experimental and biological context. Dose and delivery information may be included, but target-gene expression and cell-state variation are typically omitted, making it difficult to disentangle molecule-specific activity from context-dependent responses; (ii) chemical modifications are typically encoded as discrete identities, without capturing molecular structure or physicochemical properties; (iii) model training relies primarily on inhibition labels, underusing chemical and biophysical priors and limiting generalization to sparsely labeled targets and modification patterns.

In this paper, we propose **ASOCompass**, a context- and chemistry-aware framework for ASO activity prediction. ASOCompass first constructs a position-aligned ASO representation by combining contextualized nucleotide embeddings with molecular representations of modified monomers. It then models interactions between the ASO and its local target-RNA context through binding-aware cross-attention. Experimental conditions are captured through dose and delivery information, while biological context is modeled using transcriptome-derived representations of the target gene and cell line, adapted through learnable prototype banks. Finally, ASOCompass jointly optimizes chemical-property and sequence-derived thermodynamic objectives, encouraging its representations to capture priors related to molecular structure, stability, folding, and hybridization.

We evaluate ASOCompass on ASO Atlas (Hill et al. 2026),a large patent-derived ASO dataset, across held-out drug, target-gene, cell-line, and joint gene–cell-line settings. ASOCompass achieves the highest Spearman correlation across all four distribution shifts. On the combined test set, it achieves a Spearman correlation of 0.5970, compared with 0.5549 for OligoAI, the strongest ASO-specific baseline, corresponding to an absolute improvement of 0.0421. To assess target-specific adaptation, we exclude SOD1 and KLKB1 from supervised base-model training and fine-tune only the prediction head with limited target-specific labels. ASOCompass consistently outperforms OligoAI across few-shot label budgets and reaches Spearman correlations of 0.830 for SOD1 and 0.696 for KLKB1 with 1,024 labels.

Our main contributions are: **(i) Context-aware activity modeling**. We jointly model ASO sequence and chemistry, target-RNA context, experimental conditions, and cellular environment, enabling context-specific drug activity prediction under realistic screening conditions. **(ii) Fine-grained chemical representation**. We replace discrete modification labels with molecular representations of modified monomers, capturing fine-grained structural and physicochemical differences and their sequence-dependent effects. **(iii) Chemical and thermodynamic-guided multitask learning**. We introduce auxiliary tasks based on chemical properties and thermodynamic features to enrich activity modeling with priors on molecular conformation, target accessibility, and ASO–RNA duplex formation.

### Related Work

#### Computational ASO screening

Early computational ASO design methods ranked candidate ASO sequences based on nucleotide composition, hybridization free energy, target-site accessibility, and predicted off-target interactions (Krieg, Guga, and Stec 2003; Matveeva et al. 2003; Andersson et al. 2025; Mathews et al. 1999). RNA-folding tools such as Mfold, Sfold, and ViennaRNA further enabled estimation of secondary structure, local unpairing probabilities, and sequence-dependent binding stability (Ding, Chan, and Lawrence 2004; Zuker 2003; Lorenz et al. 2011). These approaches established useful biophysical heuristics for ASO design, but they generally relied on predefined scoring rules or simulated descriptors and did not directly model activity variation across heterogeneous experimental contexts.

With the growing availability of experimentally measured ASO datasets, recent work has increasingly focused on supervised activity prediction. ASOptimizer integrates sequence- and thermodynamics-based features with learning-guided optimization of chemical modifications (Hwang et al. 2024). OligoGym evaluates a range of sequence representations and model architectures, revealing the challenges of transferring pretrained RNA representations to chemically modified ASOs (Rotrattanadumrong and De Donno 2025). Read-across and neural approaches have likewise shown that predictive performance is strongly target-dependent and often degrades under distribution shifts (Hwang et al. 2026). Collectively, these studies underscore the importance of jointly modeling ASO sequence and chemistry. However, they generally provide limited representations of the experimental and biological contexts in which activity is measured.

Several recent methods incorporate target context or experimental metadata. eSkip-Finder uses sequence, target-site, and concentration features for exon-skipping prediction, but is restricted to a specific mechanism and narrowly defined experimental conditions (Chiba et al. 2021). OligoAI jointly encodes ASO and local target-RNA sequences while conditioning activity prediction on modification type, dose, and transfection method (Hill et al. 2026). ASOMAR combines sequence and modification encodings with thermodynamic and accessibility features for identifying effective degradation sites (Lv et al. 2026). Although these methods extend sequence-only prediction, their contextual representations remain incomplete: target-gene expression, cellular transcriptional state, and interactions among biological and experimental factors are not modeled explicitly.

#### Multimodal foundation models

Foundation models provide complementary representations for the molecular and biological components of ASO activity. RNA models can encode contextualized sequence patterns (Chen et al. 2022; Penić et al. 2025), molecular models can represent modification structure and chemical similarity (Ross et al. 2022; Zhou et al. 2023; Ji et al. 2024), and transcriptomic foundation models can capture cell-state variation (Cui et al. 2024; Kang et al. 2026). However, directly concatenating these representations is unlikely to be sufficient because they differ in scale, granularity, and pretraining objectives. ASO activity prediction therefore requires explicit alignment of position-level sequence and chemistry, interaction modeling between ASOs and target RNAs, and adaptation of transcriptomic representations to the screening task. ASOCompass is designed around these three forms of integration.

## Methods

### Problem Formulation

This paper considers context-aware prediction of ASO activity measured by inhibition percentage. Specifically, each sample is represented as *x* = (*D*, ℛ,*C*), where *D* = (*s, M*) denotes the ASO sequence and its ordered, position-aligned chemical modifications. ℛ denotes the target RNA, including the ASO-binding site and its flanking sequences. And *C* = (*d, t, g, c*) denotes the experimental and biological context, where *d, t, g*, and *c* represent the dosage, transfection method, target gene, and cell line.

As shown in Figure 2, ASOCompass jointly models the ASO drug, target RNA, and experimental and biological contexts to predict ASO inhibition:

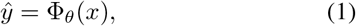

where Φ_*θ*_ denotes our proposed model. During training, we further incorporate chemical-property and thermodynamic-property prediction as auxiliary tasks to enhance representation learning. The following sections detail the model architecture and the formulation of these two auxiliary tasks.

**Figure 2:**
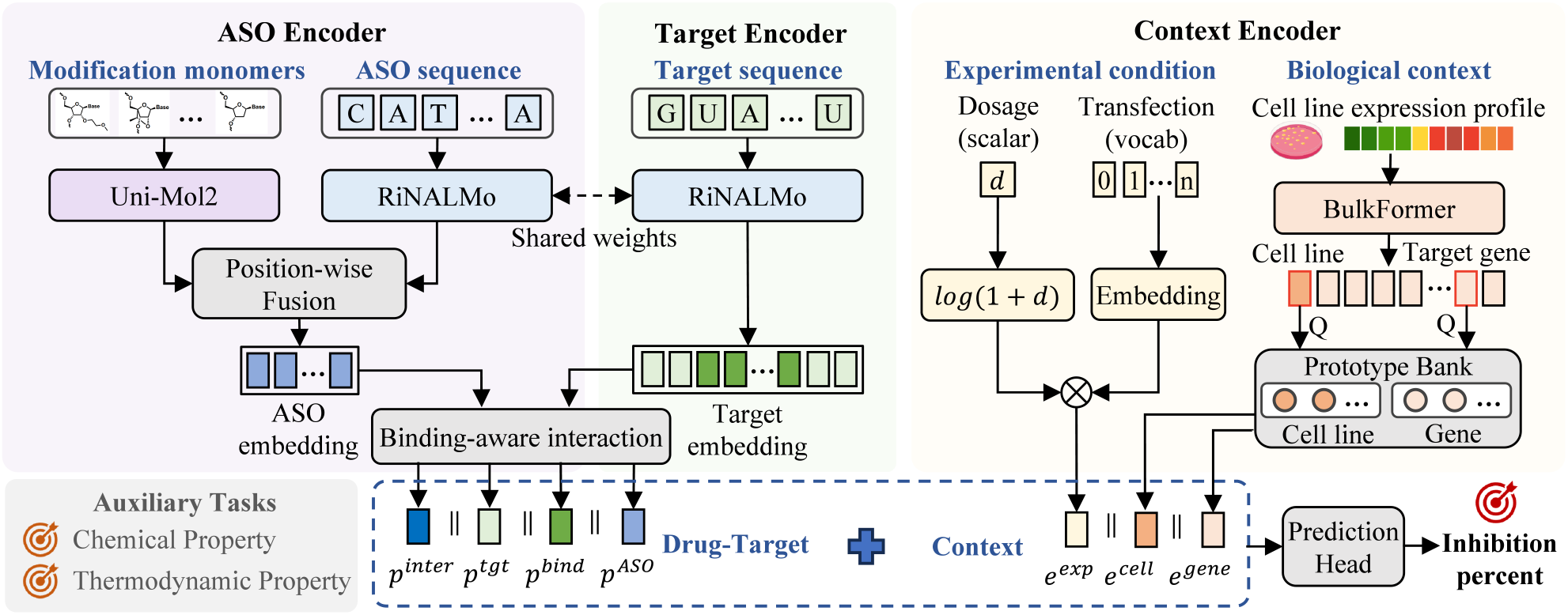
Overview of ASOCompass. ASOCompass integrates ASO sequence, target-RNA context, position-specific chemical modifications, and experimental-biological context for activity prediction. Sequence and chemical representations are fused at aligned ASO positions, while prototype-based context adapters incorporate dosage, transfection method, gene, and cell-line information. Chemical-property and thermodynamic-property auxiliary tasks provide mechanism-informed supervision.

### ASO Sequence and Chemistry Encoding

ASOs with identical nucleotide sequences can exhibit different activities due to position-specific chemical modifications. We therefore encode ASO by jointly modeling its nucleotide identity and modified-monomer structure at each position.

#### Sequence encoding

Suppose the ASO sequence *s* consists of *L* nucleotides (nt). We first use RiNALMo (Penić et al. 2025),an RNA sequence foundation model, to map *s* to contextualized nucleotide representations:

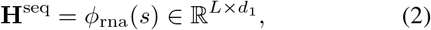

where *d*_1_ denotes the embedding dimension. For simplicity, hereafter, each *d*_*i*_ denotes the dimension of the corresponding representation.

#### Chemical encoding

For the modified-monomer sequence *M* = (*µ*_1_, …, *µ*_*L*_), where each *µ*_*i*_ is represented by a monomer-level SMILES string, we use Uni-Mol2 (Ji et al. 2024), a pretrained molecular structure encoder, to encode the chemical structure at each position:

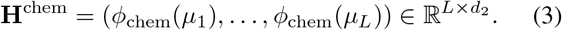

#### Position-wise sequence-chemistry fusion

We then employ a fusion module to integrate the sequence and chemical representations to obtain the final drug representation:

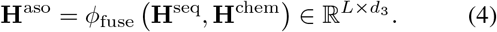

The fusion module first maps the sequence and chemical features to the same dimension using two independent gated residual projectors. It then concatenates the projected features along the feature dimension and fuses them at each position using an MLP. More details of the fusion module are provided in Appendix B. The resulting **H**^aso^ provides a position-aligned representation of the ASO, jointly modeling its sequence and chemical properties.

### Target RNA Encoding

Another factor affecting ASO efficacy is the accessibility of its target site. Although an ASO of length *L* binds to an RNA segment of the same length, accurately characterizing the accessibility of the binding site requires considering its flanking RNA context, as the local structure is influenced by the folding of the surrounding sequence.

Therefore, given the input RNA sequence ℛ of length *T* (*T > L*) that comprises the ASO-binding site and its flanking context, we use RiNALMo to encode the full sequence and then extract the binding site representations:

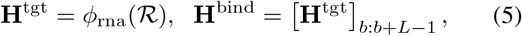

where 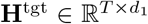, 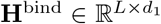, *b* denotes the starting position of the binding site in ℛ, [*·*]_*i*:*j*_ denotes the slicing operation that extracts the features from positions *i* to *j*.

### Binding-Aware ASO-Target Interaction Modeling

The representations above describe the chemically modified ASO and its target RNA separately. To explicitly model their binding interactions, we use multi-head cross-attention (MHA). Throughout this paper, MHA(**Q, K, V**) denotes multi-head attention with query, key, and value matrices **Q, K**, and **V**, respectively.

For binding-aware interaction modeling, we use the ASO representation 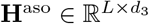 as the query and the binding-site representation 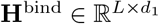 as both the key and value:

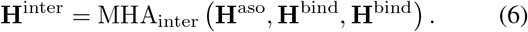

The resulting representation 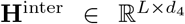 captures binding-aware target context at each ASO position and is passed to the subsequent pooling and prediction modules.

### Experimental and Biological Context Encoding

The experimental and biological context, *C* = (*d, t, g, c*), also influences the observed inhibition. The experimental context, including dosage *d* and transfection method *t*, determines ASO exposure and delivery. The biological context comprises the cell line *c*, which shapes ASO uptake, trafficking, and intracellular stability, and the target gene *g*, whose expression level affects observed inhibition.

#### Experimental context encoding

To model the experimental context (*d, t*), we first group the transfection methods *t* into four categories. Given the limited set of methods commonly used in ASO experiments: electroporation, lipofection, free uptake, and others, each transfection method category is represented by a learnable lookup embedding:

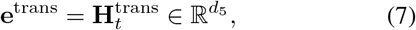

where 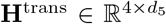 is an embedding table jointly optimized during training. Then, we incorporate the dosage *d* into the final experimental context embedding by scaling the transfection embedding:

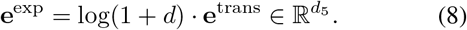

This dose-scaled representation captures the effective intracellular exposure jointly determined by dosage and transfection method.

#### Biological context encoding

To model the biological context (*g, c*), we standardize the target gene *g* using its HGNC symbol (Seal et al. 2023) and the cell line *c* using its Cellosaurus identifier (Bairoch 2018). For each cell line, we obtain a genome-wide, log-transformed gene-expression profile vector **x**_*c*_ ∈ ℝ^*G*^ from DepMap (Arafeh et al. 2025), where *G* is the number of profiled genes in this cell line. We then use a pretrained BulkFormer encoder (Kang et al. 2026) to derive transcriptome-aware representations of the cell line at each gene position:

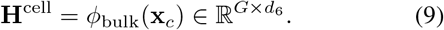

Following BulkFormer, we obtain the cell-line representation by mean-pooling across all gene-level representations, while the representation at the index corresponding to gene *g* is extracted as the target-gene representation:

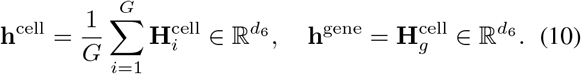

Since BulkFormer provides general-purpose gene and cell-state representations, we further adapt them to the ASO inhibition task using learnable prototypes and multi-head attention. Taking the gene representation as an example, we compute:

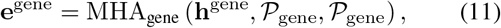

where 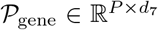 is a bank of *P* learnable prototypes, and 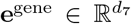 is the adapted gene representation. The cell-state representation is adapted in the same manner using a separate prototype bank *P*_cell_, yielding 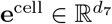.

The attention mechanism selects and combines the prototypes most relevant to each biological context, enabling the model to capture multiple aspects of biological context.

### Inhibition Prediction

After encoding all input representations, we aggregate them into fixed-length features and predict the final inhibition percentage.

#### Multi-query attention pooling

The representations of the ASO, full target context, binding site, and ASO–target interaction are variable-length feature sequences. We summarize each sequence using an independent attention-pooling module with *q* learnable queries. For feature type *a* ∈ {aso, tgt, bind, inter} and its corresponding sequence **H**, the pooling module computes

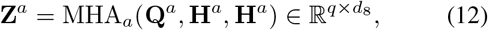

where 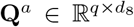 denotes the feature-specific learnable queries, and **H**^*a*^ serves as both the keys and values. The output **Z**^*a*^ consists of *q* query-specific summaries of the input sequence. These summaries are then flattened into a fixed-length representation 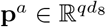.

Here, **H**^*a*^ denotes one of **H**^aso^, **H**^tgt^, **H**^bind^, and **H**^inter^. The resulting pooled representations capture distinct aspects of ASO activity: **p**^aso^ represents the ASO sequence and chemistry that govern binding and stability; **p**^tgt^ captures local RNA context related to target-site accessibility; **p**^bind^ characterizes the complementary binding region; and **p**^inter^ models ASO–target binding compatibility.

#### Final prediction

We organize the representations into two feature groups by concatenation. The ASO-target feature combines the molecular and interaction representations, while the context feature combines the experimental and biological representations:

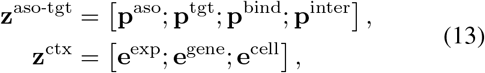

where [;] denotes feature concatenation. The two feature groups are further concatenated and passed through an MLP to predict the inhibition:

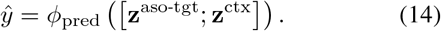

The activity loss is the mean squared error on standardized labels:

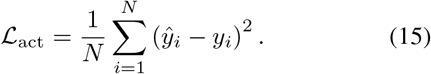

### Property-Guided Auxiliary Learning

Considering that inhibition labels provide limited supervision about the chemical and biophysical properties underlying ASO activity, we introduce two auxiliary tasks to regularize the chemical and ASO-target representations.

#### Chemical-property prediction

We construct an auxiliary dataset of ASO modified-monomer analogs from ZINC dataset (Irwin et al. 2020; Irwin and Shoichet 2005).For each molecule, we compute a set of physicochemical and quantum-chemical descriptors, as detailed in Appendix B. These chemical properties capture molecular characteristics closely associated with the stability, binding affinity, cellular uptake, and pharmacokinetic behavior of ASO therapeutics.

Given a molecular SMILES (*µ*), the model reuses the main chemical encoder from the ASO encoding branch to predict these properties:

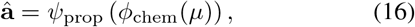

where *ψ*_prop_(*·*) is a trainable property-prediction head. We train this task using a weighted SmoothL1 loss *ℒ*_chem_.

#### Thermodynamic-property prediction

We further use ViennaRNA (Lorenz et al. 2011)to compute sequence-derived thermodynamic descriptors relevant to ASO folding and hybridization behavior. These properties are commonly considered in rational ASO design because of their close association with ASO efficacy.

Similarly, separate MLP-based thermodynamics prediction heads, *ψ*_thermo_(*·*), are adopted to map the relevant pooled ASO, target-context, and binding-region representations:

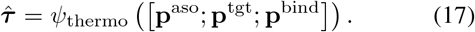

The thermodynamic prediction loss, denoted by *ℒ*_thermo_, is the mean squared error over valid descriptors.

### Training Objective

The complete training objective for ASOCompass is:

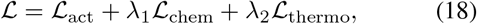

where *λ*_1_ and *λ*_2_ weight the chemical and thermodynamic losses, respectively. Further implementation details of ASOCompass are provided in Appendix C.

### Experiments

#### Experimental Setup

We evaluate ASOCompass under three settings: generalization across different distribution shifts, ablation studies of the representation modules and auxiliary objectives, and label-efficient adaptation to specialized unseen targets.

#### Datasets and preprocessing

For the main task of predicting ASO inhibition percentages, we use ASO Atlas (Hill et al. 2026), which contains 188,521 ASO experiment records extracted from 417 USPTO patents and spans 334 target genes and 31 cell lines. Each record includes the ASO sequence, position-specific chemical modifications, target gene, cell line, dosage, transfection method, target-RNA sequence with flanking context, and measured inhibition percentage. We remove records lacking target-RNA context and average replicate measurements obtained under the same ASO sequence, position-specific modification pattern, and experimental conditions, resulting in 153,174 records.

We additionally construct supervision for two auxiliary tasks. For chemical-property prediction, we compute molecular descriptors for 82,837 ZINC-derived monomer analogs. For thermodynamic-property prediction, we compute ViennaRNA-based labels from the ASO Atlas training inputs. Full details of data processing and auxiliary-task construction are provided in Appendices A and B.

#### Evaluation splits

Following the data splitting and verification procedure detailed in Appendix A, we obtain a shared training set of 104,881 records, a validation set of 19,567 records, and a test set of 28,726 records. To evaluate generalization under different distribution shifts, we partition the test set into four mutually exclusive subsets based on whether the ASO drug, target gene, and cell line appear in the training set. An ASO drug is uniquely identified by its nucleotide sequence together with its complete position-specific modification pattern. An entity is considered *seen* if it appears in the training set and *unseen* otherwise. The *drug* split evaluates generalization to unseen ASO drugs paired with seen genes and cell lines. The *gene* split evaluates unseen target genes in seen cell lines, whereas the *cell line* split evaluates unseen cell lines with seen target genes. Finally, the *gene– cell line* split evaluates the more challenging setting in which both the target gene and cell line are unseen during training. These subsets contain 12,569, 5,182, 5,008, and 5,967 records, respectively.

#### Evaluation metrics

We use the Spearman rank correlation (*ρ*) between predicted and measured inhibition percentages as the primary metric, as candidate prioritization is a central objective of ASO screening. We compute *ρ* separately for each generalization subset along with an *overall* score on the pooled set of all 28,726 test records.

#### Compared methods

We compare ASOCompass with eight classical machine learning (ML) and neural network baselines following OligoGym (Rotrattanadumrong and De Donno 2025):Linear, KNN, random forest (RF), XGBoost (XGB), MLP, CNN, GRU, and Transformer. We additionally include three ASO-specific methods: ASOMAR (Lv et al. 2026), ASORASAR (Hwang et al. 2026),and OligoAI (Hill et al. 2026). All methods were trained and evaluated on the same data partitions. Appendix D provides the feature definitions, hyperparameters, and reproduction details for every baseline.

### Overall Performance and Generalization

Figure 3(a) shows that ASOCompass achieves the strongest *overall* ranking performance among all considered methods. It obtains a Spearman correlation of 0.5970, compared with 0.5549 for the strongest baseline, OligoAI, corresponding to an absolute improvement of 0.0421 (paired permutation test, p<0.001). Classical machine-learning methods and generic neural networks achieve Spearman below 0.43, highlighting their limitations in capturing context-dependent variation in ASO activity and chemical priors underlying modifications.

**Figure 3:**
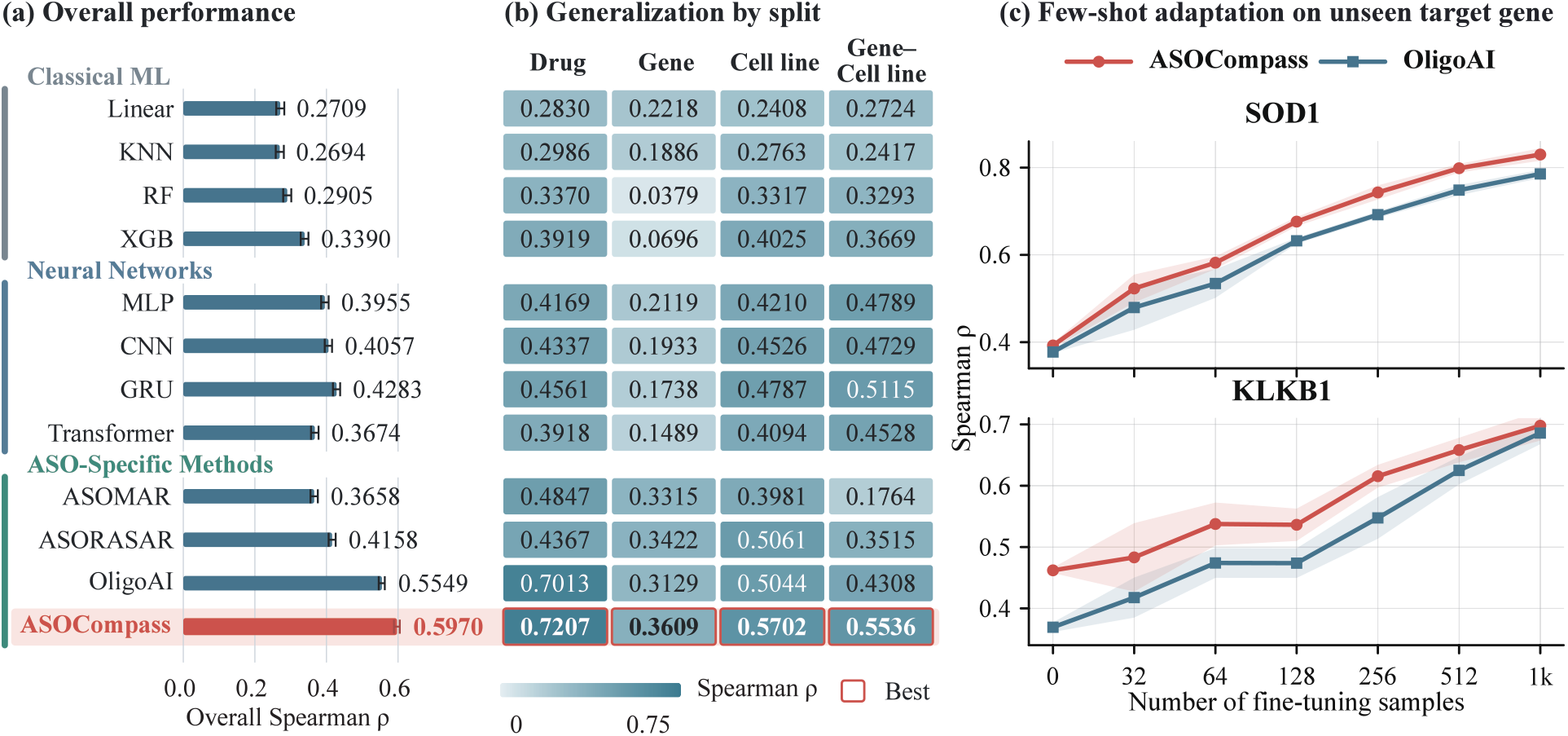
Screening performance and few-shot adaptation of ASOCompass. (a) Overall Spearman rank correlation on the complete ASO Atlas test set. (b) Spearman correlations under four distribution shifts. (c) Zero- and few-shot adaptation to the unseen SOD1 and KLKB1 targets. Lines show means and shaded bands denote standard deviation over three runs.

As shown in Figure 3(b), ASOCompass also consistently outperforms the strongest competing methods on the four distribution shifts. Specifically, it improves Spearman correlation by 0.0194 on unseen drugs, 0.0187 on unseen genes, 0.0641 on unseen cell lines, and 0.0421 on jointly unseen gene–cell line contexts. The larger improvements under cell-line and joint-context shifts suggest that integrating molecular representations with explicit biological-context modeling is particularly beneficial when generalizing to novel biological environments.

### Component Analysis

We next assess the contributions of the representation modules and auxiliary objectives through the two-stage component analysis reported in Table 1.

**Table 1:** Component analysis in terms of Spearman correlation. All auxiliary-objective variants use the same ASOCompass backbone with chemical and context encoders.

| Category | Configuration | Drug | Gene | Cell line | Gene-Cell line | Overall |
| --- | --- | --- | --- | --- | --- | --- |
| **Representation Modules** | ASOCompass (base) | 0.7013 | 0.3129 | 0.5044 | 0.4308 | 0.5549 |
|  | + Uni-Mol2 | 0.7135 | 0.3215 | 0.5286 | 0.4669 | 0.5749 |
|  | + Context | 0.7149 | 0.3540 | 0.5172 | 0.5085 | 0.5843 |
|  | ASOCompass (core) | 0.7274 | 0.3248 | 0.5274 | 0.4897 | 0.5853 |
| **Auxiliary Objectives** | + thermodynamics | 0.7184 | 0.3536 | 0.5151 | 0.5368 | 0.5892 |
|  | + chemical properties | **0.7345** | 0.3229 | 0.5280 | 0.5170 | 0.5911 |
|  | ASOCompass (full) | 0.7207 | **0.3609** | **0.5702** | **0.5536** | **0.5970** |

**Table 1:**
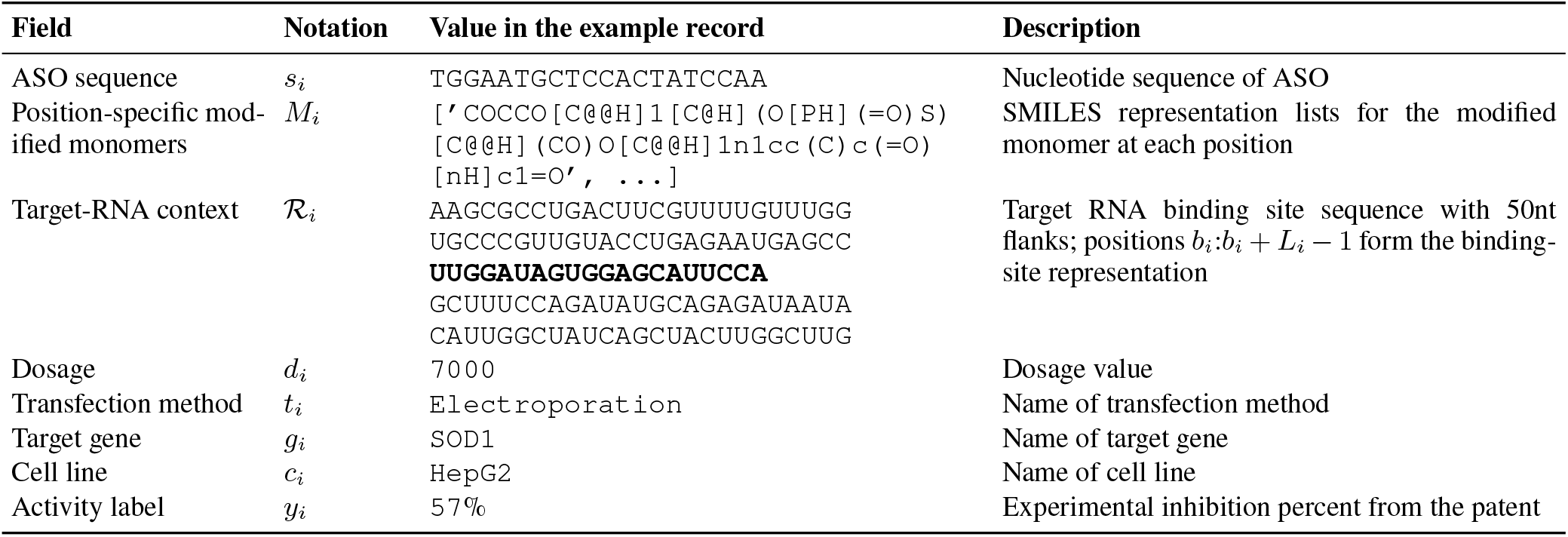
Example of one record in the ASO Atlas dataset.

We start from a plain sequence-target predictor, denoted ASOCompass (base), that consists of RiNALMo encoders for the ASO sequence and target-RNA context, categorical embeddings for position-specific sugar and backbone modifications, a dose-conditioned delivery representation, and an MLP prediction head. This model does not include molecular-structure representations, targetgene or cell-line context modules, or auxiliary objectives. Adding the Uni-Mol2 chemical encoder and biological-context modules yields the auxiliary-free model, denoted as ASOCompass (core). Jointly applying the thermodynamic and chemical-property objectives then produces ASOCompass (full). The first stage evaluates the representation modules relative to ASOCompass (base). The second stage uses ASOCompass (core) as the common reference and evaluates the two auxiliary objectives separately and jointly.

#### Biological and chemical representations improve generalization

**\**For the two individual additions to ASOCompass (base), biological context, which introduces target-gene and cell-state information, provides larger overall improvement. It raises the *overall* Spearman correlation from 0.5549 to 0.5843 and produces a substantial gain of 0.0777 in the joint *gene–cell line* generalization setting. In parallel, fine-grained molecular-structure modeling via Uni-Mol2 boosts the *overall* correlation to 0.5749, with consistent gains observed across all generalization splits. Combining these two additions produces ASOCompass (core), which achieves the strongest *overall* performance among the representation variants (0.5853) and consistently outperforms ASOCompass (base) across all four generalization splits. More component comparisons are provided in Appendix E.

#### The auxiliary objectives provide complementary supervision

When added separately to ASOCompass (core), both auxiliary objectives improve the *overall* correlation: thermodynamic supervision raises it from 0.5853 to 0.5892, while chemical-property supervision raises it to 0.5911. The chemical-property objective yields notable performance gains on unseen drugs, whereas the thermodynamic objective provides larger gains on unseen genes and joint gene– cell line contexts. Combining both objectives yields the best *overall* result (0.5970) and achieves relative improvements of 11.11%, 8.12%, and 13.05% over ASOCompass (core) on the *gene, cell line*, and joint *gene–cell line* splits, respectively, suggesting complementary benefits, particularly under biological-context shifts.

#### Chemical-property supervision improves modification-specific ranking

To examine whether chemical-property supervision improves the ability to distinguish among chemical modification designs, we construct controlled comparison groups from the drug-generalization subset. Records within a group share the nucleotide sequence, target-RNA context, dosage, delivery protocol, target gene, and cell line, and differ only in their position-specific modification patterns. We retain pairs whose measured inhibition differs by more than 15 percentage points, resulting in 129 pairs from 80 groups.

As shown in Figure 4, chemical-property supervision increases pairwise accuracy from 0.562 to 0.725 (Δ = 0.163, 95% CI [0.063, 0.271]) and Top-1 accuracy from 0.473 to 0.637 (Δ = 0.164, 95% CI [0.021, 0.299]). Top-1 accuracy measures whether the model selects the modification pattern with the highest observed inhibition within each group. The selection regret, defined as the inhibition gap between the experimentally best pattern and the model-selected pattern, decreases from 14.36 to 8.59 percentage points. These results show that chemical-property supervision specifically improves sensitivity to modification-dependent activity, rather than only increasing aggregate predictive performance.

**Figure 4:**
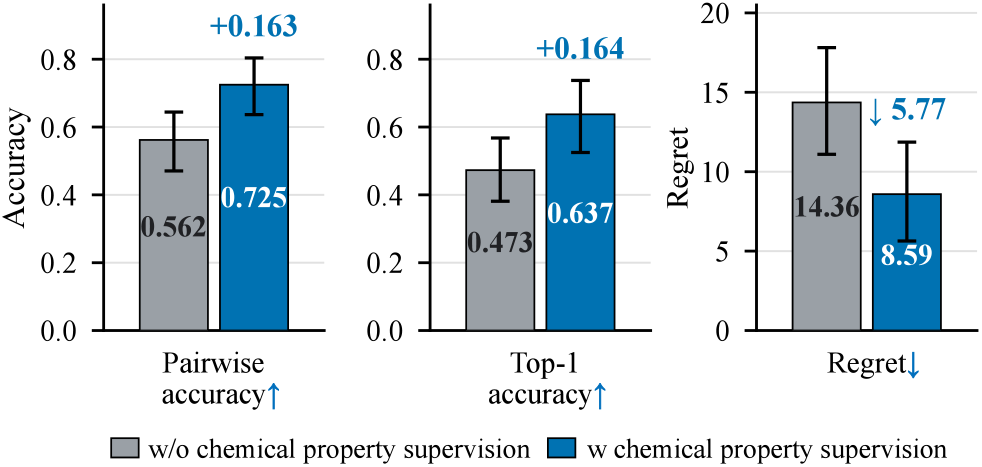
Modification-specific ranking with and without chemical-property supervision. Error bars denote bootstrap 95% confidence intervals.

Additional post-hoc analyses of attention-based ASO-pooling behavior, learned chemical representations, and the thermodynamic auxiliary representation are provided in Appendix E.

### Few-Shot Adaptation to Unseen Targets

Finally, we investigate whether ASOCompass can be adapted to a new target gene using limited target-specific activity measurements. We use SOD1 and KLKB1, two targets of FDA-approved ASO therapies with relatively abundant activity records (3,048 and 2,812, respectively). We exclude all records associated with either target from supervised base-model training. For each target, we fine-tune the prediction head using *k* ∈ {0, 32, 64, 128, 256, 512, 1024} labeled examples and evaluate it on held-out test sets to assess the adaptability of the learned representations. As shown in Figure 3(c), ASOCompass outperforms OligoAI across all annotation budgets, achieving Spearman correlations of 0.830 on SOD1 and 0.696 on KLKB1 with 1,024 labels. Notably, with only 64 KLKB1 labels, ASOCompass outperforms OligoAI trained with 128 labels (0.538 vs. 0.474), indicating that ASOCompass provides a more transferable initialization for target-specific candidate ranking. Full experimental details are provided in Appendix E.

## Conclusion

We introduce ASOCompass, a context- and chemistry-aware framework that jointly models nucleotide sequence, chemical modifications, and experimental-biological context. ASOCompass consistently improves candidate ranking across multiple distribution shifts and enables label-efficient adaptation to unseen targets. These results highlight the value of integrated molecular-context modeling for transferable ASO screening. Future work will evaluate prospective performance across broader ASO chemistries and mechanisms.

## Appendix

### A Additional Dataset Details

#### A.1 Prediction Unit and Preprocessing

Following the notation in the main paper, each activity record is represented as *x*_*i*_ = (*D*_*i*_, *ℛ*_*i*_, *C*_*i*_), where *D*_*i*_ = (*s*_*i*_, *M*_*i*_) denotes the ASO drug, with *s*_*i*_ representing its nucleotide sequence and *M*_*i*_ = (*µ*_*i*1_, …, *µ*_*iL*_) its ordered, position-aligned chemical-modification sequence represented by modified-monomer SMILES; ℛ_*i*_ denotes the target-RNA context, including the ASO-binding site and its flanking sequences; and _*i*_ = (*d*_*i*_, *t*_*i*_, *g*_*i*_, *c*_*i*_) denotes the experimental and biological context, where *d*_*i*_, *t*_*i*_, *g*_*i*_, and *c*_*i*_ represent the dosage, transfection method, target gene, and cell line, respectively. The corresponding label *y*_*i*_ is the measured inhibition percentage, clipped to [0, 100].

The activity records were derived from ASO Atlas (Hill et al. 2026). Records without a target-RNA context sequence were excluded. Repeated measurements sharing the key (*D*_*i*_, *ℛ*_*i*_, *C*_*i*_) were aggregated by averaging their inhibition percentages. The resulting dataset contained 153,174 records.

To make the prediction unit concrete, Table 1 shows one record example from the ASO Atlas dataset.

#### A.2 Generalization Split Definitions and Statistics

We evaluate all models using one shared activity training set and four mutually exclusive validation and test subsets. As shown in Table 2, the subsets are defined by the visibility of three entities in the final shared training set: the ASO drug, target gene, and cell line. Following Section A.1, the drug identity of record *i* is *u*_*i*_ = (*s*_*i*_, *M*_*i*_), namely its complete nucleotide sequence and position-specific modification pattern. Let *U*_train_, *G*_train_, and *C*_train_ denote the sets of drug identities, target genes, and cell lines occurring in the shared training set, respectively.

**Table 2:**
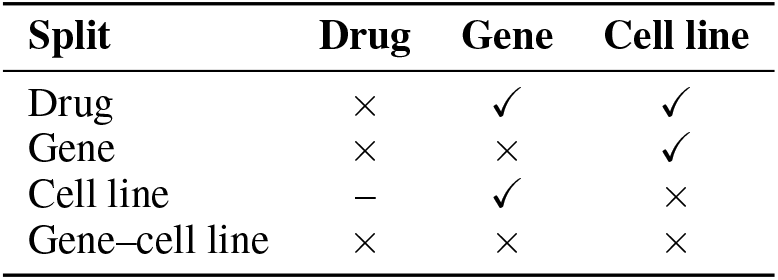
Training-set occurrence of the drug, target gene, and cell line in each evaluation split. ✓ denotes seen, *×* denotes unseen, and – denotes unconstrained.

| Split | Drug | Gene | Cell line |
| --- | --- | --- | --- |
| Drug | × | ✓ | ✓ |
| Gene | × | × | ✓ |
| Cell line | – | ✓ | × |
| Gene–cell line | × | × | × |

##### Drug generalization

This subset contains records satisfying *u*_*i*_ ∉ *U*_train_, *g*_*i*_ ∉ *G*_train_, and *c*_*i*_ ∈ *C*_train_. Thus, the complete ASO sequence–modification design is unseen, while its target gene and cell line have activity-labeled training examples.

##### Gene generalization

This subset contains records satisfying *u*_*i*_ ∉ *U*_train_, *g*_*i*_ ∉ *G*_train_, and *c*_*i*_ ∉ *C*_train_. It therefore evaluates unseen target genes and their associated ASO designs in observed cell lines.

##### Cell-line generalization

This subset contains records satisfying *g*_*i*_ ∈ *G*_train_ and *c*_*i*_ ∈*/C*_train_, and drug visibility is unconstrained. It therefore evaluates unseen cell lines for target genes observed during training.

##### Joint gene–cell-line generalization

This subset contains records satisfying *u*_*i*_ ∉ *U*_train_, *g*_*i*_ ∉ *G*_train_, and *c*_*i*_ ∉ *C*_train_. That is, the ASO, target gene, and cell line are all unseen in the training set. This subset therefore evaluates a joint shift in ASO design, target gene, and cellular context.

##### Split construction and verification

For each generalization setting, we first randomly generated candidate training, validation, and test partitions using a fixed random seed, targeting an approximate 8:1:1 ratio. We then merged them into shared partitions and assigned each validation and test record to a compatible generalization subset based on entity visibility in the shared training set. Records that matched none of the definitions were moved to the training set. After all adjustments, we recomputed entity visibility and performed a final verification of the complete partition. The resulting four evaluation subsets are mutually exclusive, and every record satisfies all constraints of its assigned subset.

##### Final partition sizes

The resulting dataset contains 104,881 training records, 19,567 validation records, and 28,726 test records. The four validation subsets and four test subsets are mutually exclusive and together form their respective composite partitions. Their record counts are reported in Table 3.

**Table 3:** Number of records in the shared training set and the mutually exclusive validation and test subsets.

| Split | Partition | Records |
| --- | --- | --- |
| Training | **Shared training set** | **104,881** |
| Validation | Drug | 12,572 |
|  | Gene | 1,560 |
|  | Cell line | 389 |
|  | Gene–cell line | 5,046 |
|  | **Total** | **19,567** |
| Test | Drug | 12,569 |
|  | Gene | 5,182 |
|  | Cell line | 5,008 |
|  | Gene–cell line | 5,967 |
|  | **Total** | **28,726** |

### B Model Components and Auxiliary Objectives

#### B.1 Position-Wise Sequence–Chemistry Fusion Module

The fusion module *ϕ*_fuse_ integrates the sequence and chemical feature sequences into a position-aligned ASO representation:

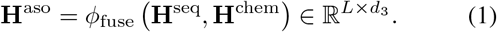

It first maps the two input feature sequences to a shared dimension using separate gated residual projectors:

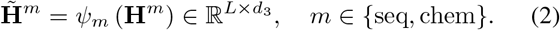

Each modality-specific projector *ψ*_*m*_ uses a gated residual architecture. Let *d*_seq_ = *d*_1_ and *d*_chem_ = *d*_2_, such that 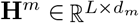. The projector is defined as

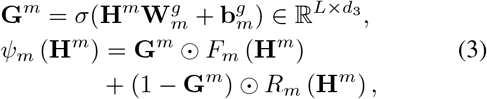

where 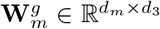 and 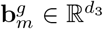 are learnable parameters, with the bias broadcast across positions. The nonlinear branch *F*_*m*_ is a two-layer position-wise MLP with layer normalization, GELU activation, and dropout, whereas *R*_*m*_ is a position-wise linear projection. The sigmoid function *σ* produces an element-wise gate **G**^*m*^ that balances the nonlinear and residual branches. The sequence and chemical projectors use the same architecture but do not share parameters.

The projected sequences are then concatenated along the feature dimension and fused using a position-wise MLP:

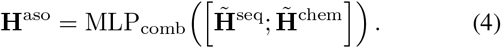

Here, the concatenated representation has dimension 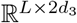, and MLP_comb_ maps it back to 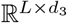. The MLP is applied independently at each ASO position.

#### B.2 Physicochemical and Thermodynamic Auxiliary Targets

ASOCompass uses two continuous auxiliary tasks. The chemical-property task is a standalone monomer-level task: a molecular SMILES representation is encoded by the shared Uni-Mol2 chemistry encoder and is used to predict 29 scalar properties. These targets comprise 24 RDKit (Landrum et al. 2025)-derived physicochemical descriptors and five semiempirical quantum-mechanical (QM) descriptors. The thermodynamic task is defined on matched ASO–target-context pairs and predicts 20 scalar descriptors calculated from the ASO sequence and its target RNA context. Each entry in Tables 4 and 5 is one-dimensional; “type” denotes the raw quantity before normalization and regression.

**Table 4:** Chemical-property auxiliary targets used by ASOCompass. Every target is a scalar (dimension 1). Units refer to the raw values before training-split normalization.

| Name | Dim. | Type | Property represented |
| --- | --- | --- | --- |
| **Charge and local electronic distribution (6 dimensions)** |  |  |  |
| estimated_charge_ph74 | 1 | Signed integer | Heuristic net charge near pH 7.4; ionization and electrostatic interaction propensity. |
| net_negative_charge_centers | 1 | Count | Number of formal or predicted deprotonated negative centers. |
| gasteiger_charge_min | 1 | Continuous | Most negative heavy-atom partial charge; strongest local electron-rich site. |
| gasteiger_charge_max | 1 | Continuous | Most positive heavy-atom partial charge; strongest local electron-poor site. |
| gasteiger_abs_charge_sum | 1 | Continuous | Total magnitude of heavy-atom partial charges; molecular polarization proxy. |
| gasteiger_p_neighbor_charge_sum | 1 | Continuous | Charge around phosphorus; local P-O/P-S/P-N backbone electronic environment. |
| **Lipophilicity, polarity, and exposed surface (6 dimensions)** |  |  |  |
| mol_logp | 1 | Continuous | Crippen octanol/water logP; lipophilicity and hydrophobic interaction propensity. |
| tpsa | 1 | Continuous ($\text{\AA}^2$) | Topological polar surface area; polarity, hydration, and hydrogen-bonding exposure. |
| labute_asa | 1 | Continuous ($\text{\AA}^2$) | Labute approximate molecular surface area; overall solvent-exposed size. |
| apolar_asa_proxy | 1 | Continuous ($\text{\AA}^2$) | $\max(0, \text{LabuteASA} - \text{TPSA})$; nonpolar exposed-surface proxy. |
| polar_surface_density | 1 | Continuous ratio | TPSA/LabuteASA; polar surface density independent of gross molecular size. |
| mol_mr | 1 | Continuous | Crippen molar refractivity; molecular polarizability and volume proxy. |
| **Size, topology, flexibility, stereochemistry, and hydrogen bonding (9 dimensions)** |  |  |  |
| exact_mw | 1 | Continuous (Da) | Exact molecular mass and overall molecular size. |
| heavy_atom_count | 1 | Count | Number of non-hydrogen atoms; molecular size and complexity. |
| rotatable_bonds | 1 | Count | Conformational flexibility and potential binding entropy cost. |
| ring_count | 1 | Count | Ring content and structural rigidity. |
| fraction_csp3 | 1 | Continuous ratio | Fraction of carbon atoms that are $sp^3$; saturation and three-dimensional character. |
| qed | 1 | Continuous [0, 1] | General small-molecule drug-likeness summary used as a low-weight baseline property. |
| chiral_center_count | 1 | Count | Assigned or unassigned stereocenters; stereochemical complexity. |
| hbd | 1 | Count | Hydrogen-bond donor capacity and hydration/recognition potential. |
| hba | 1 | Count | Hydrogen-bond acceptor capacity and hydration/recognition potential. |
| **Elemental composition and local steric bulk (3 dimensions)** |  |  |  |
| s_to_o_ratio | 1 | Continuous ratio | Sulfur-to-oxygen atom ratio; sulfur substitution and soft/polarizable character. |
| halogen_count | 1 | Count | Number of F, Cl, Br, and I atoms; halogenation, lipophilicity, and electronic effects. |
| 2prime_substituent_vdw_volume | 1 | Continuous ($\text{\AA}^3$) | Summed van der Waals volume of the C2' branch; local 2' steric bulk. |
| **GFN2-xTB quantum-chemical properties (5 dimensions)** |  |  |  |
| qm_dipole_debye | 1 | Continuous (D) | Dipole magnitude; anisotropy of the three-dimensional charge distribution. |
| qm_homo_ev | 1 | Continuous (eV) | Highest occupied molecular-orbital energy; electron-donor/oxidation tendency. |
| qm_lumo_ev | 1 | Continuous (eV) | Lowest unoccupied molecular-orbital energy; electron-acceptor/reduction tendency. |
| qm_gap_ev | 1 | Continuous (eV) | HOMO-LUMO gap; electronic softness, polarizability, and reactivity proxy. |
| qm_energy_per_heavy_atom_hartree | 1 | Continuous (Ha/atom) | Electronic energy normalized by heavy-atom count; size-adjusted energetic state. |

**Table 5:** Thermodynamic auxiliary targets used by ASOCompass. Every target is a scalar (dimension 1). ViennaRNA free energies are reported in kcal/mol before normalization; SSP denotes single-stranded probability.

| Name | Dim. | Type | Property represented |
| --- | --- | --- | --- |
| **ASO–target interaction (2 dimensions)** |  |  |  |
| dg_duplex | 1 | Continuous (kcal/mol) | ASO–target duplex free energy; more negative values indicate a more stable duplex. |
| tm_hybrid | 1 | Continuous (°C) | Nearest-neighbor RNA/DNA hybrid melting temperature; thermal duplex stability. |
| **Target-global structure (2 dimensions)** |  |  |  |
| dg_target_mfe | 1 | Continuous (kcal/mol) | MFE of the target context; global target secondary-structure stability. |
| ensemble_diversity | 1 | Continuous | Mean base-pair distance in the target ensemble; conformational heterogeneity. |
| **Target-site accessibility (10 dimensions)** |  |  |  |
| ssp_average | 1 | Continuous [0, 1] | Arithmetic mean SSP over the complete binding site; average accessibility. |
| ssp_min | 1 | Continuous [0, 1] | Minimum site SSP; the least accessible bottleneck nucleotide. |
| ssp_max | 1 | Continuous [0, 1] | Maximum site SSP; the most exposed nucleotide. |
| ssp_std | 1 | Continuous [0, 1] | Population standard deviation of site SSP; accessibility heterogeneity. |
| ssp_geomean | 1 | Continuous [0, 1] | Geometric mean site SSP; stringent accessibility summary sensitive to closed bases. |
| ssp_5p_mean | 1 | Continuous [0, 1] | Mean SSP over the first five target-site nucleotides; 5′-side accessibility. |
| ssp_3p_mean | 1 | Continuous [0, 1] | Mean SSP over the last five target-site nucleotides; 3′-side accessibility. |
| ssp_center_mean | 1 | Continuous [0, 1] | Mean SSP over the central five nucleotides; central/gap-region accessibility. |
| ssp_center_min | 1 | Continuous [0, 1] | Minimum SSP in the central five nucleotides; central accessibility bottleneck. |
| dg_opening | 1 | Continuous (kcal/mol) | Free-energy cost of exposing the target binding interval; resistance to target-site opening. |
| **ASO-only structure (4 dimensions)** |  |  |  |
| dg_aso_mfe | 1 | Continuous (kcal/mol) | MFE of the isolated ASO; self-folding stability. |
| dg_aso_opening | 1 | Continuous (kcal/mol) | Cost of forcing the complete ASO to remain unpaired; unfolding penalty before hybridization. |
| dg_aso_homodimer | 1 | Continuous (kcal/mol) | Duplex energy between two identical ASOs; self-association propensity. |
| aso_ensemble_diversity | 1 | Continuous | Mean base-pair distance in the ASO ensemble; ASO conformational heterogeneity. |
| **Composite energetics (2 dimensions)** |  |  |  |
| dg_overall_target_only | 1 | Continuous (kcal/mol) | Duplex energy plus target-opening cost; accessibility-corrected binding drive. |
| dg_overall | 1 | Continuous (kcal/mol) | Duplex energy plus target- and ASO-opening costs; net hybridization proxy. |

##### Chemical-property target calculation

###### Molecular standardization and 2D descriptors

Each SMILES was standardized with RDKit (Landrum et al. 2025) by retaining the largest fragment and applying cleanup, normalization, reionization, sanitization, and stereochemistry assignment. We used pH 7.4 as a common physiological reference for comparing monomer ionization states. Net charge was estimated with the deterministic rule:

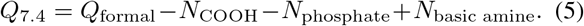

This value is a relative charge proxy rather than an exact *pK*_*a*_ or compartment-specific charge prediction. Gasteiger charges were calculated for explicit-hydrogen molecules using 24 iterations. The 2^*′*^-substituent volume was estimated from the van der Waals volume of the C2^*′*^ branch in a detected sugar-like ring and was masked when no valid context was found. The remaining RDKit descriptors and derived quantities are summarized in Table 4.

###### Semiempirical QM descriptors

For each standardized molecule, RDKit ETKDGv3 generated up to eight explicit-hydrogen conformers, pruned at a 0.5 Å RMS threshold. Conformers were relaxed and ranked with MMFF94s, with UFF used when MMFF parameters were unavailable. The lowest-energy conformer was optimized with GFN2-xTB (Bannwarth, Ehlert, and Grimme 2019) through the tblite/ASE L-BFGS interface. We retained the dipole magnitude, HOMO and LUMO energies, their energy gap, and the total electronic energy normalized by heavy-atom count.

###### Chemical-label normalization, masking, and loss

Properties with less than 5% finite coverage or zero variance were excluded; all 29 retained targets passed this criterion. Nonnegative raw targets were transformed as 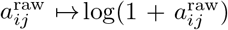, and continuous 2D and QM targets were clipped at the 0.5th and 99.5th training-set percentiles. The signed charge was not clipped. Each target was then standardized using its training-set mean and population standard deviation. Failed calculations or missing structural contexts were excluded with per-property masks. Let **a**_*i*_ and **â**_*i*_ denote the normalized target and predicted property vectors for molecule *i*, respectively. For their *j*-th entries *a*_*ij*_ and *â*_*ij*_, validity mask 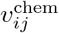, and property weight *w*_*j*_, the auxiliary loss was

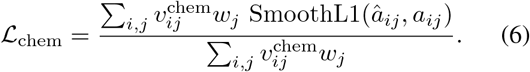

The five QM targets used *w*_*j*_ = 0.2, QED used *w*_*j*_ = 0.3, and Labute area, molar refractivity, heavy-atom count, negative-center count, phosphorus-neighbor charge, S/O ratio, and halogen count used *w*_*j*_ = 0.5; all other properties used *w*_*j*_ = 1.

###### Thermodynamic target calculation

ASO and target-context sequences were restricted to canonical A/C/G/T/U bases and converted to RNA for ViennaRNA (Lorenz et al. 2011) calculations; DNA representations were used only for melting-temperature estimation. The binding interval was taken from the site annotation when available and otherwise located by exact matching to the ASO reverse complement. ViennaRNA provided duplex free energy, minimum free energy (MFE), partition-function, and ensemble-diversity targets, while the Biopython (Cock et al. 2009) nearest-neighbor implementation with the R_DNA_NN1 parameter table provided the RNA/DNA hybrid melting temperature. Because these calculations use canonical bases and standard parameters, their outputs are physics-informed proxies rather than explicit models of MOE, cEt, phosphorothioate, or other ASO modifications.

Target-site accessibility was summarized from unpaired probabilities [ineq over the complete binding interval and its terminal and central five-nucleotide regions. Opening energy was the partition-function free-energy cost of constraining the target interval or complete ASO to remain unpaired. The two composite targets added the target-opening cost, or both target- and ASO-opening costs, to the duplex free energy. Table 5 lists the resulting targets and their physical interpretations.

For each experimental split, valid values were clipped at the 0.5th and 99.5th percentiles of the training set and standardized using the same subset. Nonfinite values, sentinel values with absolute magnitude at least 99,999, and failed calculations were masked.

###### Thermodynamic prediction heads

Equation (17) in the main paper uses *ψ*_thermo_ as compact notation for the complete thermodynamic prediction module. In the implementation, the 20 thermodynamic targets are partitioned into four groups according to the representations required for their prediction:

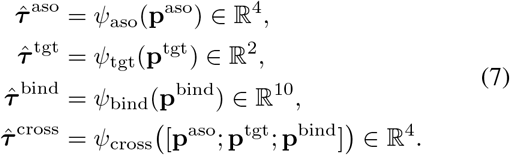

The four outputs are concatenated in a fixed order to form 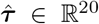. Thus, *ψ*_thermo_ in the main paper denotes this collection of group-specific MLP heads as a single vector-valued mapping.

The thermodynamic auxiliary loss was the mean squared error over valid normalized entries. Let ***τ*** _*i*_ and 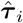 denote the normalized target and predicted thermodynamic vectors for activity record *i*, respectively. For their *j*-th entries *τ*_*ij*_ and 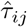 and validity mask 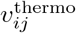, the loss was

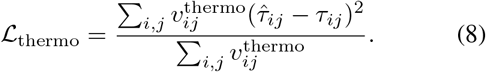

### C Implementation Details

#### C.1 Architecture and Optimization

Table 6 summarizes the principal hyperparameters that determine the reported model architecture and its optimization.

**Table 6:**
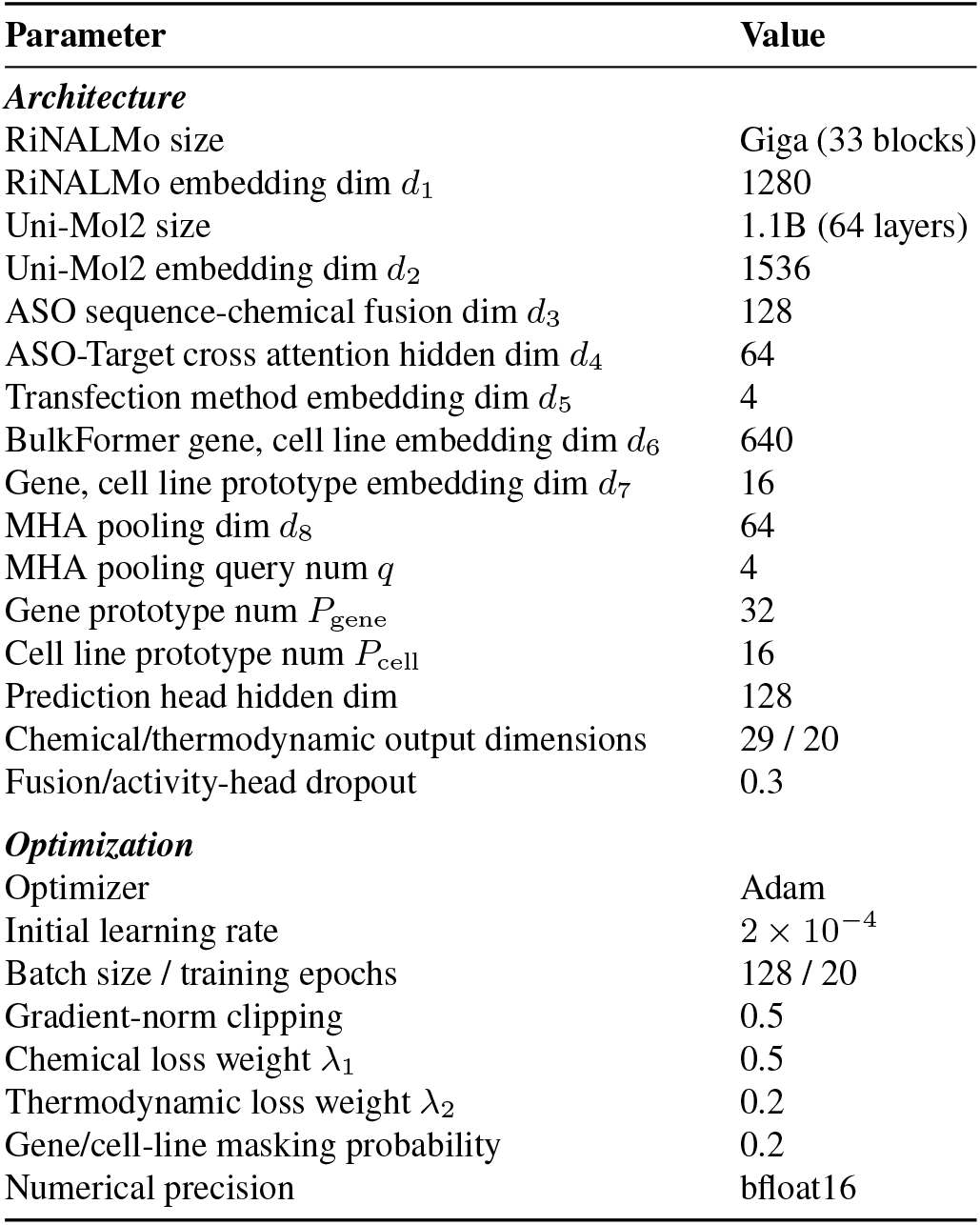
Principal architecture and optimization hyperparameters of ASOCompass.

#### C.2 Training Procedure

We trained ASOCompass for 20 epochs on a single NVIDIA A100 80 GB GPU with random seed 1 and mixed-precision bfloat16 arithmetic. The activity label was globally standardized using only the activity training split. The standalone ZINC chemical-property data were independently split 80/10/10 with the same seed. Both activity and chemical-property loaders used a batch size of 128. ViennaRNA targets were attached directly to matched examples in the activity batch.

For each joint update, the optimized objective was

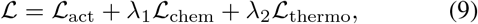

where ℒ_act_ is the mean squared error on the normalized activity label, *λ*_1_ = 0.5, and *λ*_2_ = 0.2. The masked thermodynamic and chemical losses are defined in Eqs. 8 and 6. Dropout was 0.3, and the projected gene and cell-line embeddings were independently masked with probability 0.2 during training to reduce shortcut learning. Optimization used Adam with an initial learning rate of 2 *×* 10^−4^ and gradient-norm clipping at 0.5. A step-wise linear schedule decayed the main learning rate to 10% of its initial value over the complete 20-epoch run. RiNALMo-Giga was initialized from its pretrained checkpoint, and Uni-Mol2 used the pretrained 1.1B checkpoint.

We used gradual unfreezing. At epoch 0, the pretrained RiNALMo and Uni-Mol2 backbones were frozen and only the prediction, projection, fusion, embedding, and auxiliary heads were trained. Uni-Mol2 encoder layers 62–63 were unfrozen at epoch 2. RiNALMo blocks 31–32 and its final layer normalization were unfrozen at epoch 3. At epoch 5, Uni-Mol2 layers 60–61 and RiNALMo blocks 29–30 were added; RiNALMo blocks 25–28 were added at epoch 8. Newly unfrozen parameters entered the optimizer at one tenth of the current main learning rate, while unscheduled backbone layers remained frozen.

#### C.3 Dosage and Transcriptomic Side Information

##### Dosage encoding

The merged activity table contains a single numeric dosage field, and all dosage values were reported in a consistent unit in the released dataset. For activity record *i* with delivery-protocol index *t*_*i*_ and recorded dose *d*_*i*_, the experimental-context representation is

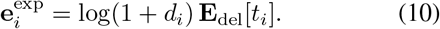

A total of 13,786 records in the dataset have missing dosage values, which are imputed with the median dosage from the training set.

##### Cell-line and target-gene context

Cell-line names were normalized using CVCL identifiers (Bairoch 2018),and bulk RNA-seq data were retrieved from the DepMap database (Arafeh et al. 2025) based on the CVCL IDs, containing log-TPM measurements for 19,215 genes. The data were fed into BulkFormer (Kang et al. 2026) to obtain cell-level and gene-level embeddings. For samples without RNA-seq data, a shared learnable missing-cell embedding was used. If the cell line was covered but the target gene could not be indexed, a separate shared learnable missing-gene embedding was used.

### D Comparison Methods

#### D.1 OligoGym

##### OligoGym implementation

Our general machine-learning baselines follow the implementations in OligoGym (Rotrattanadumrong and De Donno 2025). OligoGym represents an oligonucleotide by its nucleobase sequence and position-aligned sugar and backbone modifications. The three categorical components are independently one-hot encoded and concatenated. For an ASO of length *L*, the resulting representation is

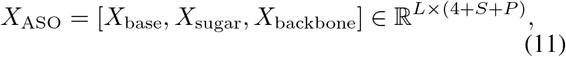

where *S* and *P* denote the numbers of sugar and phosphate categories, respectively. OligoGym benchmarks Linear, KNN, RF, XGB, MLP, CNN, GRU, and Transformer models. Positional one-hot features are flattened for Linear, KNN, RF, XGB, and MLP, whereas CNN, GRU, and Transformer retain the sequence axis. The statistical models use scikit-learn or XGBoost, and the neural models use PyTorch Lightning. OligoGym searches predefined grids over model depth, hidden dimensionality, convolution and pooling choices, and dropout. Its neural models are optimized with Adam for at most 100 epochs, with early-stopping patience of five epochs.

##### Settings in our reproduction

We retrained these eight model classes on our data following the above model definitions. All models used the full position-wise one-hot representation in Eq. 11, including the nucleobase, sugar, and backbone channels. Category vocabularies were fitted on the training set only; unseen validation or test categories were mapped to all-zero vectors. Sequences were right-padded with zeros to the maximum ASO length in each data table. The tabular models and MLP received flattened vectors, while the three sequence models received the unflattened matrices. Table 7 lists the fixed configurations used in our experiments.

**Table 7:** Fixed configurations of the OligoGym-based baselines in our reproduction.

| Model | Configuration |
| --- | --- |
| Linear | Ridge regression with $\alpha = 1.0$. |
| KNN | $k = 5$, Euclidean distance, and uniform neighbor weights. |
| RF | 100 trees with maximum depth 10. |
| XGB | 100 trees with maximum depth 10 and the squared-error objective. |
| MLP | One hidden layer of width 128, ReLU activation, and dropout 0.25. |
| CNN | A single one-dimensional convolutional layer with 64 channels, kernel width 5, same padding, and global max pooling. |
| GRU | One recurrent layer with hidden dimensionality 64; the final hidden state is used for prediction. |
| Transformer | Embedding dimensionality 128, two layers, four attention heads, feedforward dimensionality 128, dropout 0.25, and mean pooling. |

**Table 8:** Extended component analysis measured by Spearman correlation. Each row adds the indicated chemical or biological-context component to ASOCompass(base); auxiliary objectives are excluded. The best result in each column is shown in bold.

| Configuration | Overall | Drug | Gene | Cell line | Gene-cell line |
| --- | --- | --- | --- | --- | --- |
| ASOCompass (base) | 0.5549 | 0.7013 | 0.3129 | 0.5044 | 0.4308 |
| + Uni-Mol2 (frozen) | 0.5745 | 0.7058 | **0.3623** | 0.5201 | 0.4682 |
| + Uni-Mol2 (fine-tuned) | 0.5749 | 0.7135 | 0.3215 | **0.5286** | 0.4669 |
| + Context (learnable embedding) | 0.5570 | 0.6875 | 0.3282 | 0.5127 | 0.4525 |
| + Context (prototype pool) | **0.5843** | **0.7149** | 0.3540 | 0.5172 | **0.5085** |

All eight models performed single-target regression of ASO inhibition percentage using molecular features only; cell line, target gene, transfection method, and dosage were excluded. RF and XGB used random seed 42. Neural models were trained with mean squared error and Adam using a learning rate of 10^−3^, weight decay of 10^−2^, batch size 256, and at most 100 epochs. The data splitting strategy is the same as that used in the main experiment. Training stopped if validation loss did not improve for five consecutive epochs, and the checkpoint with the lowest validation loss was retained.

#### D.2 OligoAI

##### Method overview

The ASO Atlas study introduces OligoAI, a multimodal regression model for RNase H-mediated ASO efficacy prediction (Hill et al. 2026). OligoAI applies pretrained RiNALMo to separately encode the ASO sequence and its local target pre-mRNA context. At each ASO position, learned embeddings of the sugar modification (MOE, cEt, or DNA) and backbone linkage (PS or PO) are combined with the sequence representation before pooling. The pooled ASO and target representations are then concatenated with a transfection-method embedding scaled by log(1 + *d*), where *d* is the dosage, and an MLP predicts the inhibition percent.

##### Settings in our reproduction

We reimplemented OligoAI on the processed ASO Atlas records used in our main experiments. We retained the published RiNALMo-giga sequence encoders, the *±*50-nt target-context window, the discrete sugar and backbone embeddings, the dosage transformation, and the original fusion and prediction architecture. Transfection protocols were mapped to electroporation, free uptake, lipofection, or other, following the original implementation. To isolate the published OligoAI feature set, this baseline did not use molecular-structure encodings from Uni-Mol2, gene or cell-line representations, prototype adapters, or the chemical and thermodynamic auxiliary objectives introduced in ASOCompass.

For direct comparison with the other methods, we replaced OligoAI’s original data split with the same training, validation, and test partitions used in our main experiments; all other implementation and training settings followed the original study.

#### D.3 ASO-RASAR

##### Method overview

ASO-RASAR (Hwang et al. 2026) is a read-across sequence–activity relationship framework that integrates a global quantitative structure-activity relationship (QSAR) model, target-specific expert models, and nearest-neighbor similarity features using a second-stage meta-model. The global component learns activity patterns shared across genes, while the expert predictions and similarity features transfer information from data-rich targets to targets with limited or unseen training data. We adapted the released classification pipeline to predict continuous inhibition percentage and ranked ASOs according to their predicted values.

##### Settings in our reproduction

We retained the original feature specification, including 47 genomic, energy, structure, and off-target descriptors, sequence 4-mer counts, and positional nucleotide one-hot features. The data partition was kept consistent with our main experiments. Within the training partition only, five-fold grouped cross-fitting was used to generate out-of-fold global-QSAR predictions for fitting the second-stage meta-model. We trained 16 random-forest expert models (300 trees; maximum depth 20) and a random-forest meta-model (100 trees; maximum depth 10). The high- and low-inhibition similarity pools were separated using the target-specific 80th percentile estimated from the training data, and all training references were retained. We used random seed 0 and computed Spearman correlation on the test set.

#### D.4 ASOMAR

##### Method overview

ASOMAR (Lv et al. 2026)is a two-branch neural network that combines the ASO sequence with experimentally and thermodynamically derived descriptors. The sequence branch applies a one-dimensional convolutional neural network to the one-hot encoded ASO sequence, while the descriptor branch uses an MLP to encode chemical modification, experimental concentration, ASO folding and self-hybridization, ASO–target hybridization, melting temperature, and target accessibility. The representations produced by the two branches are concatenated and processed by a shared prediction head.

##### Settings in our reproduction

The original ASOMAR model performs binary classification of highly and weakly effective ASOs. Since our task is to predict the continuous inhibition percentage, we retained its two-branch architecture but replaced the two-class softmax layer with a single neuron with linear activation. The model was trained using mean squared error and the Adam optimizer with a learning rate of 1 *×* 10^−3^ and a batch size of 128. All other settings remain consistent with the original paper.

### E Additional Experimental Results

#### E.1 Extended Component Analysis

To complement the component analysis in the main paper, we further isolate two implementation choices: adaptation of the pretrained chemical backbone and the parameterization of biological context. Each variant adds only the indicated component to ASOCompass(base) and is evaluated using the same training and generalization protocol as the main experiments. The fine-tuned Uni-Mol2 variant is adapted during activity training, whereas its frozen counterpart keeps the pretrained encoder fixed. The two context variants compare direct learnable context embeddings with the prototype-pool adapter used by ASOCompass.

##### Controlled fine-tuning provides modest task adaptation

Fine-tuning Uni-Mol2 yields an overall correlation comparable to that of the frozen encoder (0.5749 versus 0.5745), indicating that the pretrained chemical representations already transfer effectively to ASO activity prediction. Nevertheless, fine-tuning improves the drug and cell-line results by 0.0077 and 0.0085, respectively. The gain on the drug split is particularly relevant because this setting evaluates transfer to unseen ASO designs and therefore directly tests the utility of the chemical representation. We adopt the fine-tuned configuration to retain the overall performance of the frozen encoder while allowing additional task-specific alignment on new molecular candidates. The gradual, low-learning-rate schedule in Appendix C.2 restricts this adaptation to the upper encoder layers, balancing task alignment with preservation of the pretrained chemical representation.

##### Prototype pools make biological context transferable

Replacing direct learnable context embeddings with the prototype-pool adapter raises the overall correlation from 0.5570 to 0.5843 and improves all four generalization splits. The largest gain is obtained for jointly unseen gene–cell-line contexts, where the correlation increases by 0.0560 (0.4525 to 0.5085). Moreover, direct embeddings provide only a small overall gain over ASOCompass(base) (0.0021), whereas prototype adaptation improves it by 0.0294. These results are consistent with the prototype pool sharing task-relevant structure across biological contexts instead of representing each context as an unconstrained embedding, thereby supporting transfer to unseen gene and cell-line combinations.

#### E.2 Few-Shot Adaptation to Unseen Targets

##### Targets and data isolation

We use SOD1 and KLKB1 to evaluate supervised adaptation to previously unseen target genes. The two targets contain 3,048 and 2,812 records, respectively. In this experiment, all records associated with either target are excluded from supervised base-model training. Consequently, before few-shot adaptation, neither model has observed target-specific inhibition labels for these genes. We compare our model, ASOCompass, with OligoAI, which achieved the best *overall* performance among the baselines.

##### Episode construction

We construct three independent adaptation episodes for each target. In repeat *r*, all records associated with the target are randomly permuted using split seed *s*_*r*_ 1, 2, 3 and divided equally into an adaptation pool and a test set. This produces 1,524 adaptation and 1,524 test records for SOD1, and 1,406 adaptation and 1,406 test records for KLKB1. The test set is fixed across all annotation budgets within a repeat, but is reconstructed independently across repeats.

For a given repeat, the *k*-shot labeled records for model fine-tuning consist of the first *k* records in the shuffled adaptation pool. The resulting fine-tuning sets are nested: if *k*_1_ *< k*_2_, then the *k*_1_-shot set is a strict subset of the *k*_2_-shot set. This construction reduces sampling-induced variation when comparing adjacent annotation budgets. The complete few-shot adaptation run evaluates

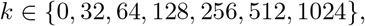

The *k* = 0 condition directly evaluates the target-independent checkpoint without parameter updates. The same split seeds and resulting support/query partitions are used for both models.

##### Adaptation procedure

Each (target gene, repeat, *k*) run is independently initialized from the corresponding model’s fixed pretrained checkpoint. During adaptation, only the prediction head is optimized. Each model is optimized for 20 epochs with a batch size of 16. We use Adam with an initial learning rate of 2 *×* 10^−4^ and a step-wise linear decay to 2 *×* 10^−5^ over the complete adaptation run. The objective is mean squared error on globally standardized inhibition labels. The normalization statistics are fitted exclusively on the original base-training split and are held fixed for every target, repeat, and annotation budget.

##### Evaluation and aggregation

After adaptation, we evaluate the final-epoch model on the test set associated with the same repeat. We compute Spearman’s rank correlation between predicted and measured inhibition values over the complete test set. Reported results are the mean and population standard deviation over the three independently seeded repeats.

##### Results

ASOCompass outperforms OligoAI at every reported annotation budget. With 1,024 labeled target-specific records, ASOCompass obtains Spearman correlations of 0.830 *±* 0.014 on SOD1 and 0.696 *±* 0.021 on KLKB1, compared with 0.786 *±* 0.009 and 0.686 *±* 0.018, respectively, for OligoAI. Cross-budget comparisons further demonstrate improved label efficiency. With only 64 KLKB1 labels, ASO-Compass reaches 0.538 0.035, exceeding the 0.474±0.024 obtained by OligoAI with 128 labels. Similarly, ASOCompass with 512 SOD1 labels achieves 0.799 *±* 0.010, surpassing OligoAI with 1,024 labels (0.786 *±* 0.009). All comparisons are computed from the held-out test sets and constitute a separate post-training adaptation analysis.

#### E.3 Position-Wise Analysis of ASO Pooling Attention

The ASO-specific learnable-query pooling module compresses the position-resolved representation **H**^aso^ into the fixed-length representation **p**^aso^. ASOs are short synthetic oligonucleotides designed to bind complementary target RNAs. RNase H gapmer ASOs contain a central DNA gap that supports RNase H1-mediated cleavage of the target RNA, with chemically modified wings at the 5^*′*^ and 3^*′*^ ends that improve binding affinity and stability. Gapmer design therefore commonly involves varying the types and positional patterns of modifications within these wings. This architecture motivates us to examine whether the pooling module attends differently to gap and wing positions.

We perform two complementary analyses to examine how the pooling module distributes attention across these regions. The first compares the aggregate positional profiles of 16-nt and 20-nt ASOs, testing whether a broad wing–gap pattern is consistent across ASO lengths. The second fixes the ASO length at 16 nt and stratifies records by their exact sugar-modification patterns, testing whether the positional profile also varies with the chemical layout.

##### Length-level positional profiles

We analyzed the ASO-pooling attention weights to determine which nucleotide positions were preferentially retained by the ASO-specific representation branch. This analysis included 12,221 16-nt ASOs and 14,470 20-nt ASOs from the held-out test set. Attention assigned to special tokens was excluded, and the remaining weights were renormalized over nucleotide positions and averaged across attention heads and pooling queries. As shown in Figure 1, the model exhibited a clear wing-dominant attention profile for both ASO lengths. In 16-nt ASOs, positions 1-3 and 14-16 received substantially higher attention than the uniform expectation of 1/16, whereas the central positions received markedly lower weights. A similar pattern was observed for 20-nt ASOs: positions 1-5 and 16-20 were consistently enriched relative to the uniform expectation of 1/20, while positions 6-15 were strongly down-weighted. These positional profiles closely correspond to the terminally modified wings and central DNA gap of the dominant gapmer architectures in the test set. Because the ASO-pooling module operates on representations integrating nucleotide identity and per-position chemical features, the results suggest that the model preferentially retains information from chemically modified terminal regions when constructing its ASO-level representation.

**Figure 1:**
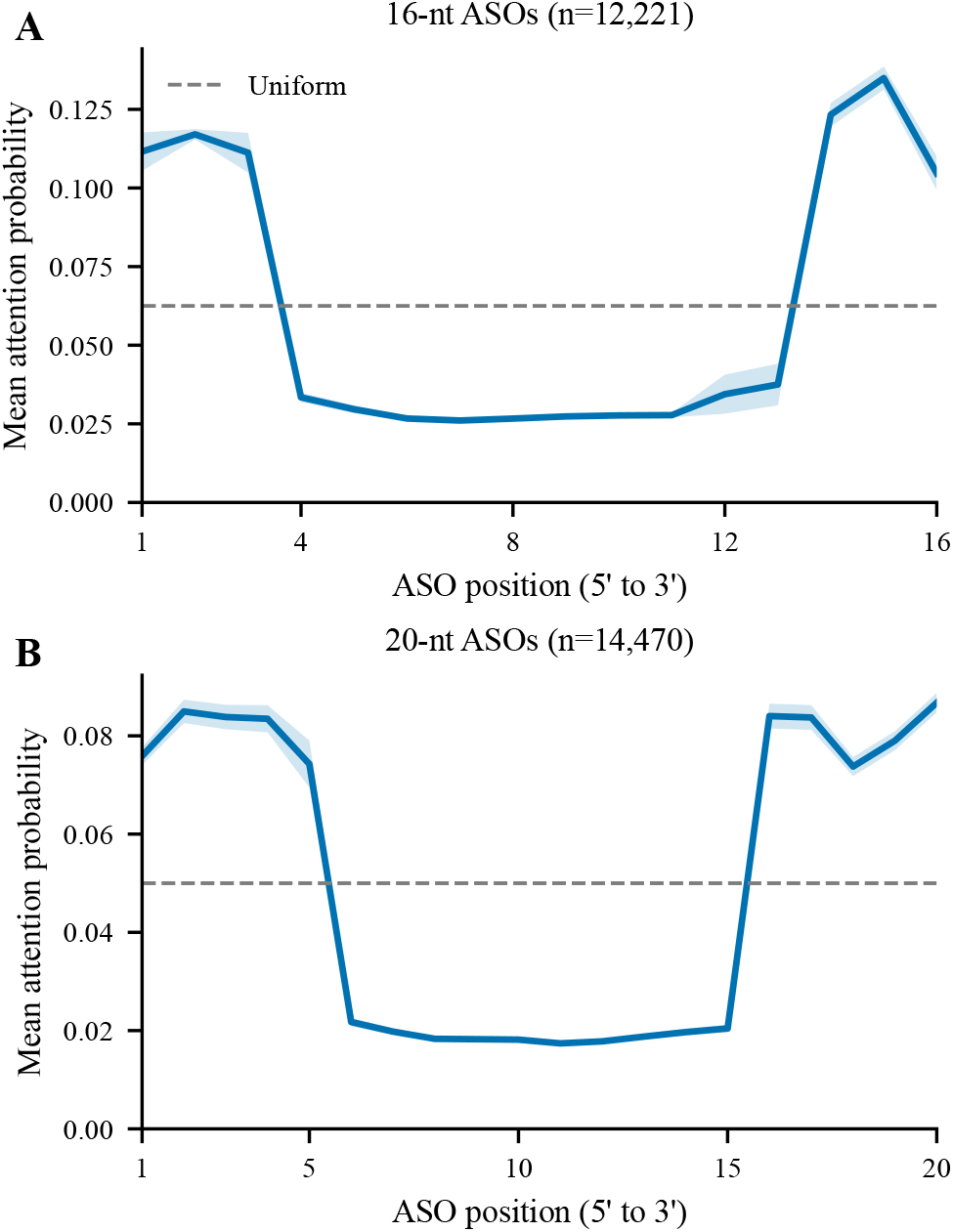
ASO-pooling attention profiles for 16-nt and 20-nt ASOs. Solid lines show mean attention probabilities over the held-out test set, and shaded regions indicate 95% confidence intervals. Dashed lines denote uniform attention. Both lengths exhibit enriched attention at the terminal wings and reduced attention within the central DNA gap, indicating that the ASO-specific pooling module preferentially aggregates features from terminally modified regions.

##### Modification-pattern-specific profiles

The length-level analysis averages ASOs with different chemical layouts and therefore cannot distinguish a general terminal-position preference from variation associated with specific modification patterns. To examine this distinction, we restrict the analysis to 16-nt ASOs and group the held-out records by their exact position-wise sugar-modification patterns. We retain the six most frequent patterns, denoted S1–S6, which together contain 10,733 records. We process the ASO-pooling attention weights as above and average them across records within each pattern. For visualization, each position’s mean attention is divided by the uniform expectation of 1*/*16, such that a value of 1 denotes uniform attention.

Figure 2 shows that the positional profiles are not identical across chemical designs. A localized comparison between S1 and S2 further illustrates this pattern dependence. Relative to the common 3–10–3 cEt gapmer pattern represented by S1, S2 replaces the cEt residues at positions 1 and 16 with MOE while retaining the remaining sugar-modification layout. Both substituted terminal positions exhibit pronounced attention enrichment, consistent with the pooling module responding to position-specific changes in sugar chemistry rather than only reproducing a fixed terminal preference. More generally, wings containing cEt- or MOE-modified terminal regions receive greater attention at those positions and lower attention within their central DNA regions. By contrast, S6, which has an unmodified all-DNA sugar pattern, exhibits a substantially flatter profile without a pronounced wing–gap separation. Together, the two analyses provide complementary evidence: the length-level comparison identifies a wing-dominant preference that persists across 16-nt and 20-nt ASOs, whereas the pattern-stratified comparison shows that attention allocation also varies with the position-specific sugar-modification layout.

**Figure 2:**
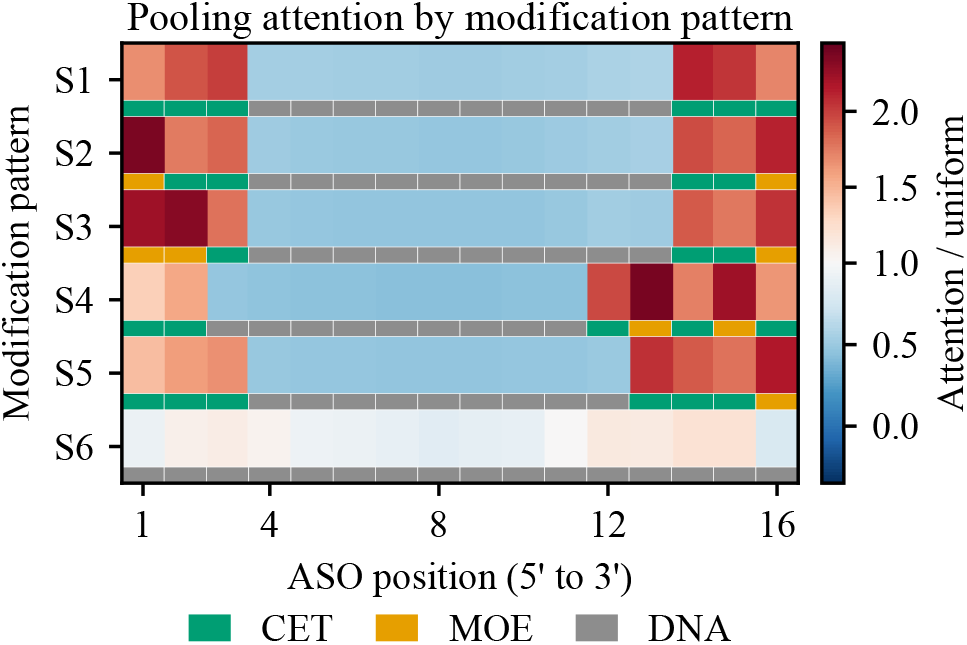
Sugar-modification-pattern-specific ASO-pooling attention for the six most frequent 16-nt patterns. Each row corresponds to one pattern, and the colored strip below the row identifies the sugar chemistry at each position. Heatmap values report attention relative to the uniform expectation of 1*/*16; values above 1 indicate enrichment.

#### E.4 Chemical Representation Analysis

To examine whether the chemistry branch learns chemically meaningful representations rather than merely encoding nucleotide identity, we extracted the representations of all 39 modification monomers in the ASO Atlas dataset after the trained chemical projection layer. After *ℓ*_2_ normalization, each monomer was used to retrieve its *k* nearest neighbors (*k* ∈ {1, 3, 5}) by cosine similarity. We measured the fraction of retrieved neighbors sharing the same complete sugar-backbone modification, sugar modification, backbone modification, or nucleobase, and compared it with class-frequency-matched random expectations using 10,000 label permutations. If the learned space captures modification chemistry, monomers sharing sugar or backbone modifications should cluster together and yield retrieval precision above the random baseline; conversely, enrichment by nucleobase would suggest that the representations are dominated by nucleotide identity.

**Figure 3:**
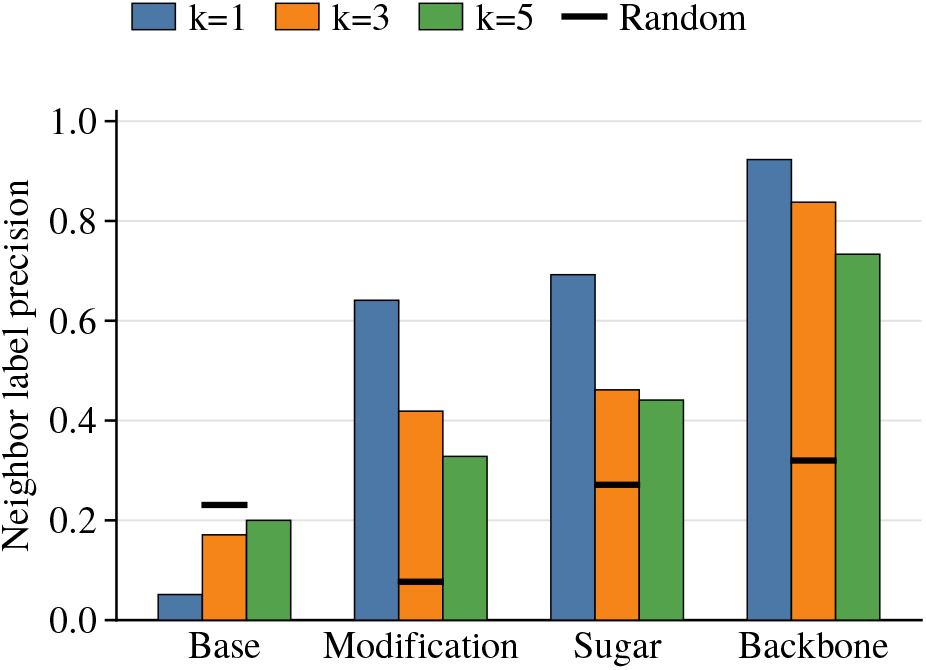
kNN analysis of learned monomer representations. Bars show same-label precision among the top-*k* cosine neighbors; black ticks indicate random expectations. The representations capture modification chemistry, particularly backbone structure, rather than base identity.

As shown in Figure 3, the nearest-neighbor space was strongly organized by modification chemistry: at *k* = 1, the retrieval precision reached 64.1% for the complete modification label, 69.2% for sugar chemistry, and 92.3% for backbone chemistry, compared with random expectations of 7.7%, 27.1%, and 32.0%, respectively. These enrichments remained significant for all evaluated values of *k* (permutation *p* ≤ 6 × 10^−4^). In contrast, nucleobase identity was not enriched among neighboring monomers (5.1–20.0% versus a 23.1% random expectation). Together, these results provide representation-level evidence that ASOCompass organizes monomers according to fine-grained modification chemistry, particularly backbone and sugar chemistry, rather than simply grouping them by nucleobase identity.

#### E.5 Activity Association Profile of the Thermodynamic Proxy

We investigated whether the thermodynamic proxy preserves the relationships between ViennaRNA descriptors and experimentally measured ASO activity. This analysis was performed on the same held-out ASO Atlas test set used in the main experiments, containing 28,726 records. For each of the 20 thermodynamic descriptors, we separately calculated its Spearman association with the inhibition-percentage label using (i) the descriptor directly computed with ViennaRNA and (ii) the corresponding value predicted by the thermodynamic auxiliary head.

Figure 4 compares the resulting feature–activity association profiles. Each point represents one thermodynamic descriptor, with its horizontal and vertical coordinates denoting the inhibition associations obtained from the directly computed and predicted values, respectively. The predicted descriptors closely reproduce the activity-association pattern of the computed descriptors: the Spearman correlation between the two 20-dimensional association profiles reaches 0.943. Moreover, 19 of the 20 descriptors have consistent association signs, and the mean absolute difference between the corresponding correlation coefficients is only 0.024. A permutation test over the descriptor identities further confirms that this agreement is unlikely to occur by chance (*p <* 10^−4^). The consistency is observed across descriptor groups, including duplex interaction, target structure and accessibility, ASO folding, and composite energy features. These results indicate that the auxiliary head preserves which thermodynamic factors are positively or negatively associated with measured inhibition, rather than merely approximating their numerical values.

**Figure 4:**
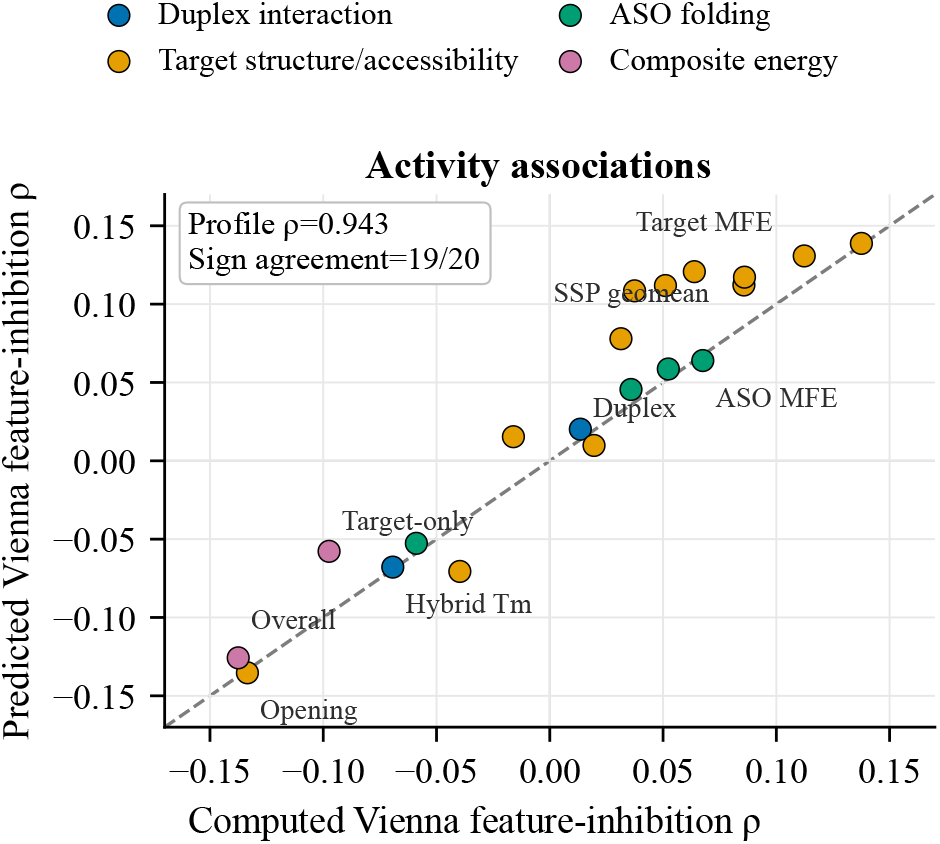
Agreement between the activity associations of directly computed and model-predicted ViennaRNA descriptors. Each point represents one of 20 thermodynamic descriptors and is positioned according to its Spearman correlation with measured inhibition percentage. The dashed diagonal denotes identical associations for the computed and predicted descriptors. Colors indicate the corresponding thermodynamic feature groups.

#### E.6 Activity Readout of the Thermodynamic Auxiliary Representation

We examine whether the ViennaRNA auxiliary task learns thermodynamic representations relevant to experimentally measured inhibition. This post-hoc diagnostic uses the same held-out ASO Atlas test set as the primary evaluation, comprising 28,726 records: 12,569 in the *drug* split, 5,182 in the *gene* split, 5,008 in the *cell line* split, and 5,967 in the *gene–cell-line* split. It does not alter the training or reported predictions of the primary ASOCompass model.

For every record, we collect the 20 descriptors computed directly with ViennaRNA and the corresponding 20 values predicted by the trained thermodynamic auxiliary head. Each descriptor set is then used separately as input to a linear Ridge readout and a nonlinear gradient-boosted readout for measured inhibition percentage. Readout performance is evaluated using five-fold cross-validation. This experiment measures activity alignment rather than the physical accuracy of the head-predicted thermodynamic quantities.

As shown in Fig. 5, the head-predicted descriptors yield stronger out-of-fold inhibition readouts than the directly computed descriptors. For Ridge, the overall Spearman correlation increases from 0.172 to 0.208 (Δ = +0.036, 95% CI [0.017, 0.052]); for gradient boosting, it increases from 0.228 to 0.273 (Δ = +0.045, 95% CI [0.028, 0.063]). Improvements are positive on all four generalization splits, ranging from +0.003 to +0.075, and are largest for unseen genes and jointly unseen gene–cell-line contexts. The auxiliary objective therefore appears to produce thermodynamically informed features that are more closely aligned with measured inhibition, rather than merely reproducing isolated computed quantities.

**Figure 5:**
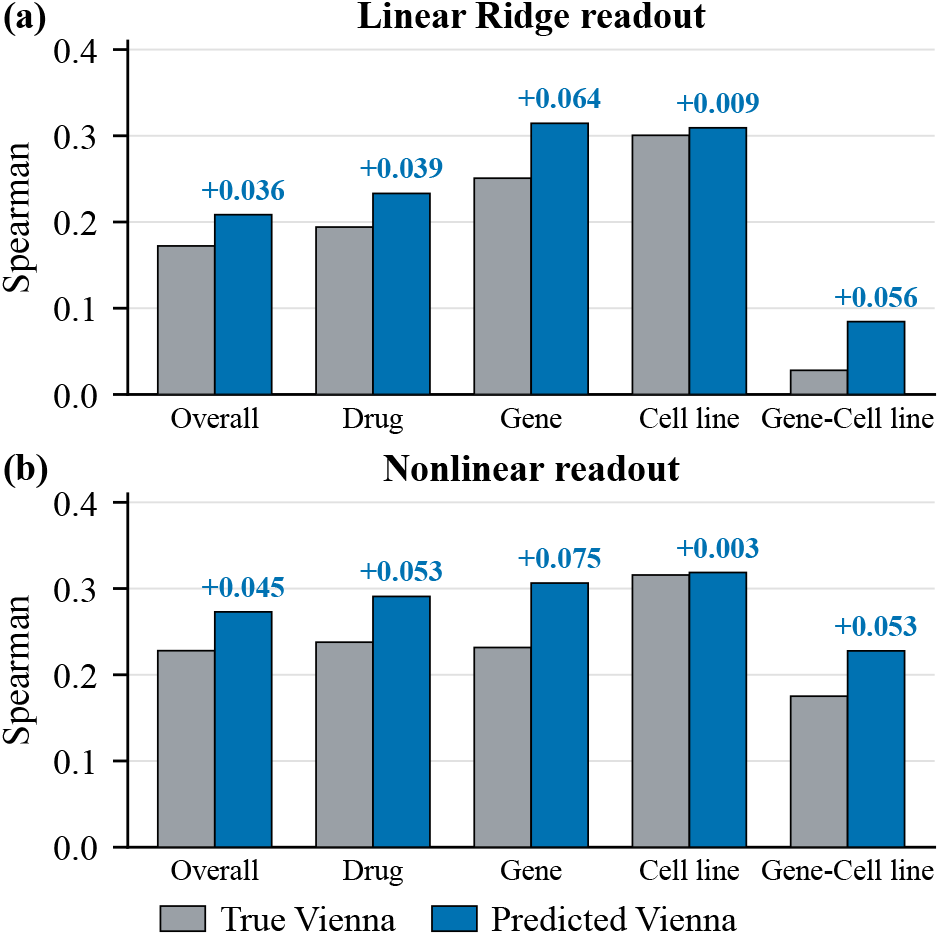
Activity prediction from directly computed and head-predicted ViennaRNA features using Ridge (a) and gradient boosting (b). Bars show out-of-fold Spearman correlations under five-fold cross-validation; labels give the absolute gains from predicted features.

### F Statistical Analysis

Statistical analyses were performed using per-case predictions from the held-out test set. The 95% confidence intervals for overall Spearman correlations were estimated using 5,000 nonparametric bootstrap resamples of paired observations and predictions, with percentile-based confidence limits. Differences between models within each generalization split were assessed using two-sided paired permutation tests on matched cases, with prediction ranks exchanged within each pair over 99,999 permutations. Monte Carlo-corrected *p*-values were reported.

